# Monotherapy cancer drug-blind response prediction is limited to intraclass generalization

**DOI:** 10.1101/2025.06.16.659838

**Authors:** William G. Herbert, Nicholas Chia, Paul A. Jensen, Marina RS Walther-Antonio

**Affiliations:** Graduate School of Biomedical Sciences, Mayo Clinic, Rochester, MN, USA; Department of Obstetrics and Gynecology, Mayo Clinic, Rochester, MN, USA; Department of Surgery, Mayo Clinic, Rochester, MN, USA; Microbiome Program, Center for Individualized Medicine, Mayo Clinic, Rochester, MN, USA; Computing, Environment, and Life Sciences, Argonne National Laboratory, Lemont, IL, USA; Department of Biomedical Engineering, University of Michigan, Ann Arbor, MI, USA; Department of Chemical Engineering, University of Michigan, Ann Arbor, MI, USA

## Abstract

Monotherapy cancer drug response prediction (DRP) models predict the response of a cell line to a given drug. Analyzing these models’ performance includes assessing their ability to predict the response of cell lines to new drugs, i.e. drugs that are not in the training set. Drug-blind prediction displays greatly diminished performance or outright failure across a wide range of model architectures and different large pharmacogenomic datasets. Drug-blind failure is hypothesized to be caused by the relatively limited set of drugs present in these datasets. The time and cost associated with further cell line experiments is significant, and it is impossible to predict beforehand how much data would be enough to overcome drug-blind failure. We must first define how current data contributes to drug-blind failure before attempting to remedy drug-blind failure with further data collection. In this work, we quantify the extent to which drug-blind generalizability relies on mechanistic overlap of drugs between training and testing splits. We first identify that the majority of mixed set DRP model performance can be attributed to drug overfitting, likely inhibiting generalization and preventing accurate analysis. Then, by specifically probing the drug-blind ability of models, we reveal the sources of generalizable drug features are confined to shared mechanisms of action and related pathways. Furthermore, we observed that, for certain mechanisms, we can significantly improve performance by limiting the training of models to a single mechanism compared to training on all drugs simultaneously. We conclude that drug-blind performance is a poor benchmark for DRP as it does not describe model behavior, it describes dataset behavior. Our investigation displays that deep learning models trained on large, monotherapy cell line panels can more accurately describe mechanism of action of drugs rather than their advertised connection to broader cancer biology.

**Author summary:** In this paper, we characterize the feature space of cancer drug-blind prediction. To understand the efficacy of a novel cancer drug it has never seen before (drug-blind), a model must be able to accurately compare this drug to all drugs it saw during training. These relationships between cancer drugs, the feature space, must be described well enough that this is possible. We believe that these relationships are poorly defined because cancer DRP models always display reduced performance in a drug-blind context. For the first time, we quantified the limits of generalization in a drug-blind setting. We showed that drug-blind generalization describes mechanistic relationships among drugs during model training. We also outlined new criteria with which to judge the drug-blind ability of a model. Failure of drug-blind prediction is an oft overlooked shortcoming in cancer DRP with potentially damaging downstream implications. We hope to show drug-blind ability of these models in a new light to guide others towards more pertinent tasks in cancer deep learning.

## Introduction

Five year cancer survival rates have almost doubled in the past five decades, a remarkable sign that our understanding of the disease is improving [1]. An inherent side effect of this is that as we develop successful treatments, we are left with increasingly complex cancer cases that require an increasingly intricate examination of cancer biology [2]. The application of artificial intelligence (AI) has been touted as the solution to unlocking an understanding of cancer that would allow us to overcome this [3, 4]. The translational function DRP models seek to recapitulate is the identification of a compound that will be effective in treating a specific type of cancer.

Over 100 different cancer DRP models have been published utilizing large pharmacogenomic datasets and attempting to translate advances in AI to improve our understanding of cancer treatment [4]. Recent work has focused on ensuring the robustness of these models by developing frameworks for reusability and standardized comparison [5, 6]. While this increased rigor for is useful for quantifying relative performance gains, the question remains - what are we actually learning about cancer treatment when training these models on large drug response panels?

To answer this question, we can look toward methods for training and quantifying performance of cancer DRP models. Architecture-focused approaches hypothesize that different deep learning model designs will extract different, and perhaps better, information from cancer cell line drug response data. The majority of these models are trained on the same datasets and response metrics [4]. Very few of these models compare to previously published models, instead comparing to out-of-the-box machine learning baselines [6]. When the reusability of these models was assessed, it was determined that for many of them it was difficult to obtain the original claimed performance with hurdles including minimal detail on preprocessing or hyperparameter optimization [6]. This creates challenges in DRP model benchmarking due to inconsistency in analysis and lack of robust training details.

Rather than designing different model architectures, we can examine how training data drives DRP model behavior. There are a variety of different large monotherapy pharmacogenomic cancer cell line panels used in cancer DRP training [7]. Early work examined the general scaling laws applicable to these datasets [8]. This is important for guiding further collection and experimentation. Different datasets also display different levels of efficacy for prediction among one another, showing that not all cancer DRP datasets are created equal. [9] These works set the stage for examination of cancer drug response generalizability.

We can evaluate model performance itself in different data-centric manners. The most common form of analysis is in the form of mixed set testing, where all drugs and cell lines may appear repeatedly in training and testing. This is the least robust form of model evaluation. More translatable analyses are cancer-blind and drug-blind splits. These are models where the sets of unique cell lines and drugs, respectively, are disjoint in the training and test set.

Cancer-blind and drug-blind analysis both describe the ability of a model to generalize. Models are traditionally capable of cancer-blind prediction, but performance worsens significantly in a drug-blind setting and some models fail completely [4]. This failure is incredibly concerning given that drug features should be generalizable in order to be transferable to new treatments or more broadly applicable to patients. The relevance of cancer cell lines themselves with respect to patients itself is already questionable [10]. If we are to determine the difference in drug generalization in a patient setting versus a cell line setting, we must first begin to understand drug generalization in a cell line setting alone.

When determining the capability of an AI model or dataset to perform a task, it is important to thoroughly probe potential sources of failure [11, 12]. This is particularly true when the downstream applications are sensitive, such as in healthcare. The shortcomings of large pharmacogenomic datasets themselves are well documented [13–16], but little has been done to explain causes of drug-blind failure. Our primary objective, then, is to examine exactly what occurs during model training to drive drug-blind failure.

It is essential to place our work in the context of prior studies that perform drug-focused analysis of cancer DRP performance. Model performance has been shown to improve when trained on all drugs at once rather than on each drug individually. [17]. This work indicates that there is beneficial drug-to-drug information which appears to cluster by mechanism of action. It is also clear that there is disparate information that can be learned from each drug and that some drugs are more difficult to learn than others [18]. Taken together with the prior study, there appears to be both constructive information among drugs tempered by a heterogeneity among learnable features. It was found that experimental noise can damage learned chemical features, but binary representation of drug response restored some performance in drug-blind models [19]. Furthermore, specific aggregations of results can be used to provide a more reliable evaluation of drug-blind performance [20]. In one of the only efforts to directly address drug-blind failure, performance was improved using multi-objective optimization to prevent training from skewing performance to specific drugs by enhancing the extraction of global drug response features [21].

In this work, we sought to understand the specifics of what makes drug-blind prediction difficult. What this amounted to was the careful examination of how designing different training and testing splits impacts what a model learns and generalizes to. Previous approaches in drug discovery attempt to “learn” the optimal composition of training data to maximize performance with as few training examples as possible [22]. There are also approaches in quantitative structure activity relationship modelling for rational selection of train/test splits [23]. In cancer drug response prediction, the overlap between chemical structures in the training and testing set presents a difficult confounding factor when analyzing model performance. Indeed, prior works examining drug-blind prediction discuss this as a limitation, even recommending that one might consider tuning the training and testing sets to particular model applications [20, 21]

Rather than developing an application for addressing drug-blind failure, we sought to fully characterize drug-blind limitations. We find that, as hypothesized, drug-blind failure occurs because the feature space describing cancer drugs is incomplete. While individual mechanisms are well defined, global features describing cancer drugs are nonexistent. Little to no relevant information exists in our models for a previously unseen drug if it does not fall into these well defined mechanisms. Different model architectures have not and will not solve drug-blind failure because this information does not exist to be extracted from the actual data they are trained on. Moreover, it is difficult to purposefully fill in these gaps in the feature space. Further characterization can reveal relationships among pathways, but it is not feasible to predict these relationships in entirely novel mechanisms. Mechanism focused model training provides a potential solution by focusing learning on well-defined regions of the feature space. We conclude that future DRP work must prioritize appropriate depth and variety in a mechanism specific setting rather than breadth across global drug response.

The key contributions of this work are as follows:

1. We quantify how information sharing occurs across drugs in a drug-blind setting. We find that information sharing occurs primarily among drugs with the same mechanism of action and, to a lesser extent, drugs with related mechanisms. Certain kinase inhibitors with broader affinity benefit from shared information across many classes.
2. We show that models identify drugs with similar mechanisms during training using only cell line responses. Models reinforce mechanistic information, if it is available, rather than overfitting to individual drugs.
3. We observe confining model training to specific mechanisms of action better resolves individual drug-cell line pairs by decreasing drug to drug similarity. These models can learn behavior of drugs on specific cell lines rather than collapsing to learning mechanims of action.

## Results

### Permutation of response values reveals dependence on drug distributions

Drug response values were permuted either within drug (intradrug shuffle) or within cell lines (intracell shuffle) to determine whether models are learning a true drug-cell line connection (Fig. 1). Intradrug shuffling and intracell shuffling identify the dependence of a model on the specific pairing of a drug with a cell line or a cell line with a drug, respectively. We hypothesized that any impacts of permutation on performance are driven by the heterogeneity and distribution of learned responses rather than to learned label noise as in prior studies [13, 17]. One-hot encoding removes all biological information and replaces it with a binary for the cell line present. This creates a similar disconnect from cell line to drug as intradrug shuffling, but retains true cell-drug pairings.

**Fig 1.**
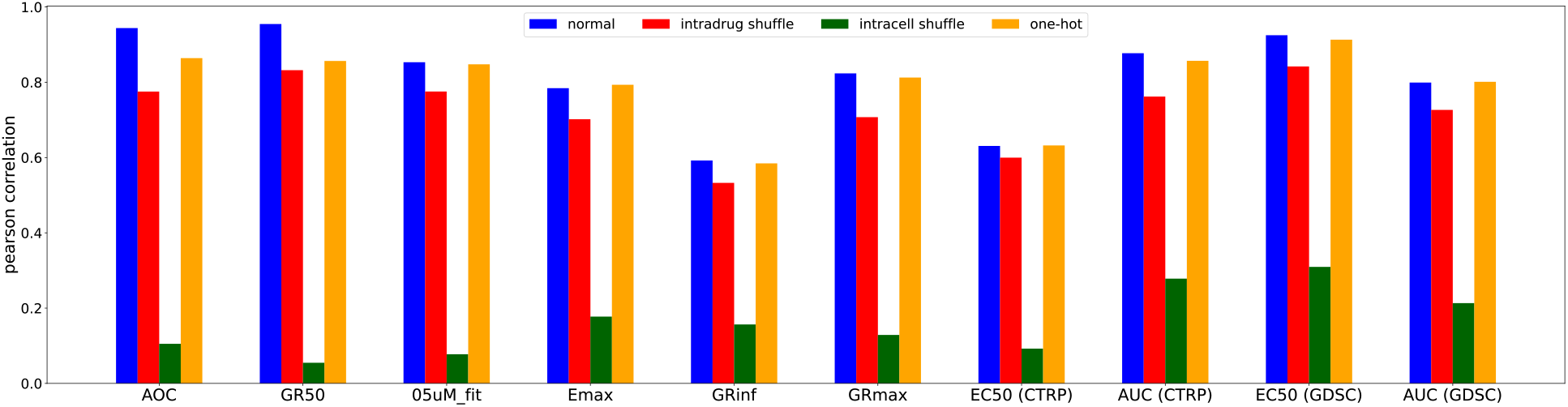
Permutation of response data across cell lines and drugs. We examine model performance across all included datasets and metrics for 3 different input conditions and a control. Response metrics are denoted on the x-axis, with source datasets included for repeats.

We expected that randomly shuffling the data would remove information about specific cell-line/drug pairings and reduce performance. Intradrug shuffling decreased performance by only 10–15% for all metrics across all three datasets, while intracell shuffling deteriorated performance by 60–90%. This suggests learning the overall response distribution of a particular drug, not any specific drug-biology connections, determines up to 90% of DRP model performance. Learning specific drug-biology connections is essential to exrapolating to unseen drugs and biological measurements taken from tumors; however, our models are overfitting to individual drugs without concern for cell line features.

### Quantifying the impact of dataset diversity on performance

We next sought to examine the limits of learning the underlying distributions of drug response. In particular, we examined the impact on performance of limiting the amount of cell lines that describe a drug’s behavior. We predicted that as a drug is tested on more cell lines, the better the drug’s overall response distribution will be defined. In actuality, we saw there is no appreciable drop off until around 40 cell line examples when decreasing the number of cell line examples in the training set per drug (Fig. 2A). Overall, this indicates that efforts to broaden the scope of cancer drug response datasets should focus on the addition of unique drugs rather than repeatedly testing a drug on different cell lines. This result confirmed prior work examining the utility of increasing the scope of cell line experiments across datasets [9].

**Fig 2.**
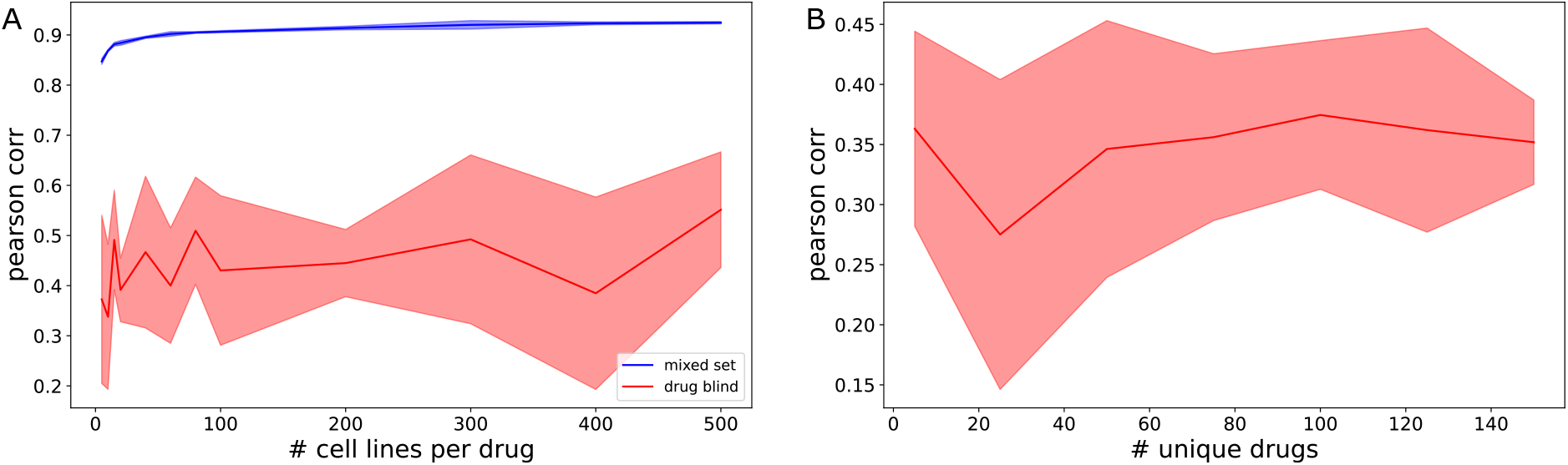
Drug-blind performance exhibits greater variance compared to mixed set testing. (A) DRP model performance when increasing the number of cell lines per drug between mixed set and drug blind conditions and (B) when changing the number of unique cell lines in training set under drug-blind conditions. Shaded regions indicate standard deviation across five replicates with varying train/val/test splits. Increasing both number of cell lines per drug and unique drugs in training set does not improve performance, but variance greatly increases from mixed set to drug blind testing. Interestingly, the variance of results obtained increased drastically between mixed set and drug-blind testing.

Previous work has hypothesized that increasing the amount of unique drugs in a dataset would improve drug-blind performance since model performance increases with dataset size both within and across datasets [4]. As far as we are aware, this has never been shown empirically with respect to drug-blind performance. Limiting the amount of unique drugs present in the training set does not decrease average model performance, but variance of performance decreases as more drugs are added to the training set (Fig. 2B). This decrease in variance is caused by increased overlap between the chemical properties of the training and test set when more unique drugs are added during training.

### Training set composition predicts drug-blind performance

We determined the association between the drugs contained in the training set and overall performance by randomly varying drugs present during training across a set of models (Fig. 3A). The pearson correlation for all models trained varies between 0.517 and 0.169 with a standard deviation of .079. Given such variability, we fit a linear model with input drugs as features and model performance as targets (Fig. 3B). The elastic net model showed a clear connection between drugs in the training set and overall performance even with a small sample size of experiments. While we used a constant sized training set, it is important to consider that as a higher proportion of total drugs are included during training, overall performance variance will decrease, as in Fig. 2B. Creating a predictive model of performance requires that a smaller subset of total unique drugs be used because of this.

**Fig 3.**
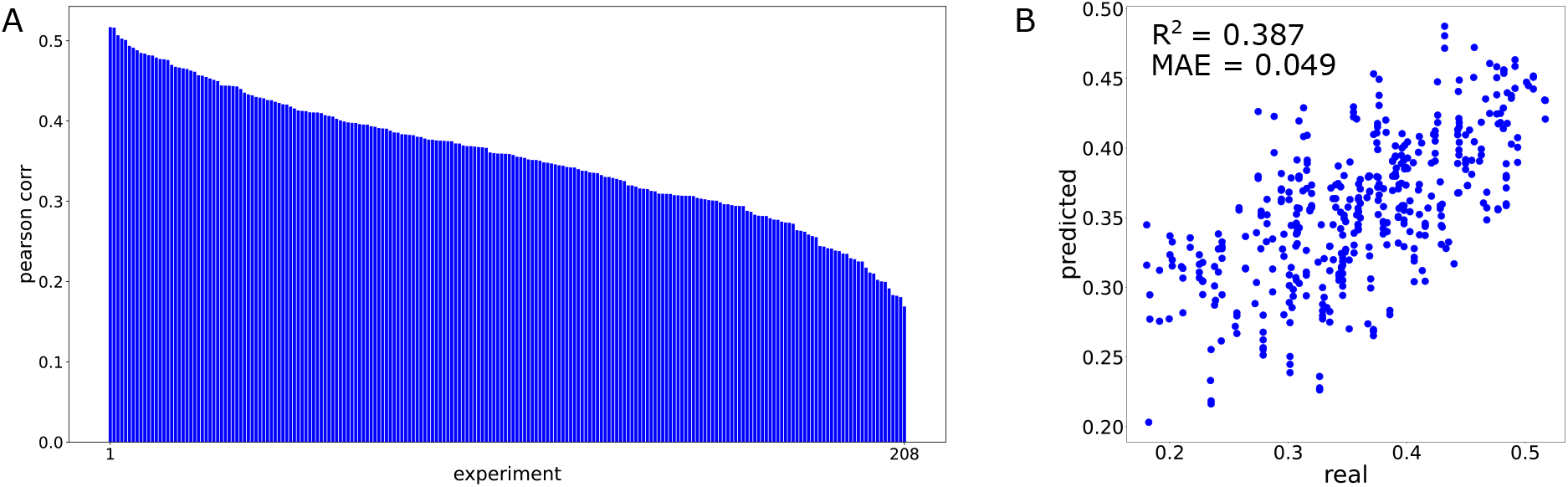
Drugs in training set predict overall performance of drug-blind test set. (A) Individual performance across 208 drug-blind models with varying training sets and constant test set. Each bar is a single model. (B) Predictions made by elastic net model fit to model performance using training set composition. Each point is a single model.

### Drug-blind generalization is limited to functional relationships

We then extended the experiment from the prior section to an approach where training set composition is similarly varied, but the test set is composed of all drugs available in the dataset. Therefore, drugs are be drug-blind in some models but not in others. A linear model was trained with input drugs in the training set as the features and accuracy for an individual drug across all experiments as the targets. Coefficients for drugs as predictors are hierarchically clustered to create a heatmap of these associations (Fig. 4). More positive coefficients indicate that the presence of a particular drug in the training set improves our prediction of response accuracy of a particular drug in the test set, while negative coefficients indicate a loss of predictability.

**Fig 4.**
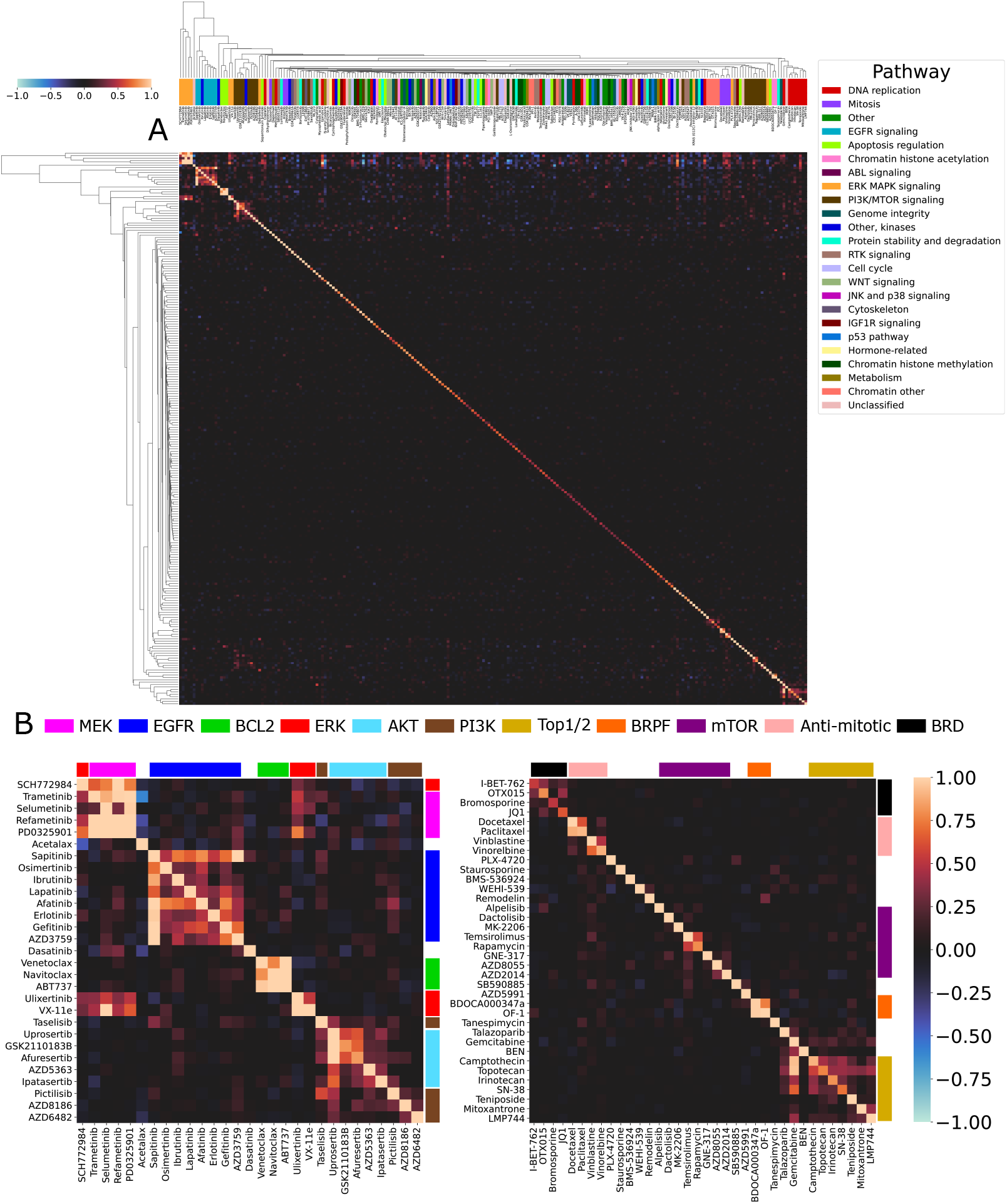
Clustering of coefficients reveals intraclass drug-blind generalization. (A) Drugs as features (y-axis) are trained to predict individual test set drug performance (x-axis) using elastic net across 1,641 trained DRP models. Dendrogram is created using agglomerative clustering of coefficients for each individual drug. The presence of the diagonal in this clustering dendrogram displays that most predictive power for a drug’s accuracy is when itself is in the training set (non-drug-blind). Performance prediction outside of self-association corresponds to mechanisms of action. Cluster color indicates the GDSC supplied pathway annotation. Plotted values are capped at [-1 1] to increase contrast and improve visualization. Best viewed zoomed in. (B) Magnified view of core cancer drug target pathways identified by drug-blind generalization. Coefficients from elastic net model fit to drugs as predictors (y-axis) of the performance of targets in the test set (x-axis). Drugs are color coded according to their advertised specific mechanism of action rather than GDSC supplied pathway annotations.

As expected, the diagonal of the coefficient heatmap contains the largest values i.e. the best predictor for the performance of a drug in the test set is itself. This is the non-drug-blind case. For 224 out of 230 drugs (97%), the strongest predictor of performance is the non-drug-blind case. This gives some insight into the extreme performance drop in drug-blind experiments. During mixed set testing, drugs are highly dependent on their own examples for prediction. Furthermore, we see that coefficients in the center of the heatmap are incredibly sparse. Not only is the strongest predictor of performance almost always the drug itself, most drugs don’t have any other predictors of performance. This indicates that for the majority of drugs, there exists little to no relevant information learned by the model being shared from drug to drug.

Outside of self-association, we see squares of shared coefficients. These clusters reveal themselves to represent different cancer drug mechanisms of action (Fig. 4). The bulk of identified clusters are part of the RAF/MEK/ERK and PI3K/AKT/mTOR pathways. These pathways are central to cell proliferation and are both commonly mutated and targeted in many different cancers [24, 25]. Interestingly, the coefficients determining drug predictive power extend across these pathways, which we term class-to-class relationships (Fig. 5C). The presence of an ERK or MEK inhibitor in the training set improves the prediction of the other in the test set. ERK is commonly represented as being immediately downstream from MEK in cellular signaling pathways.

**Fig 5.**
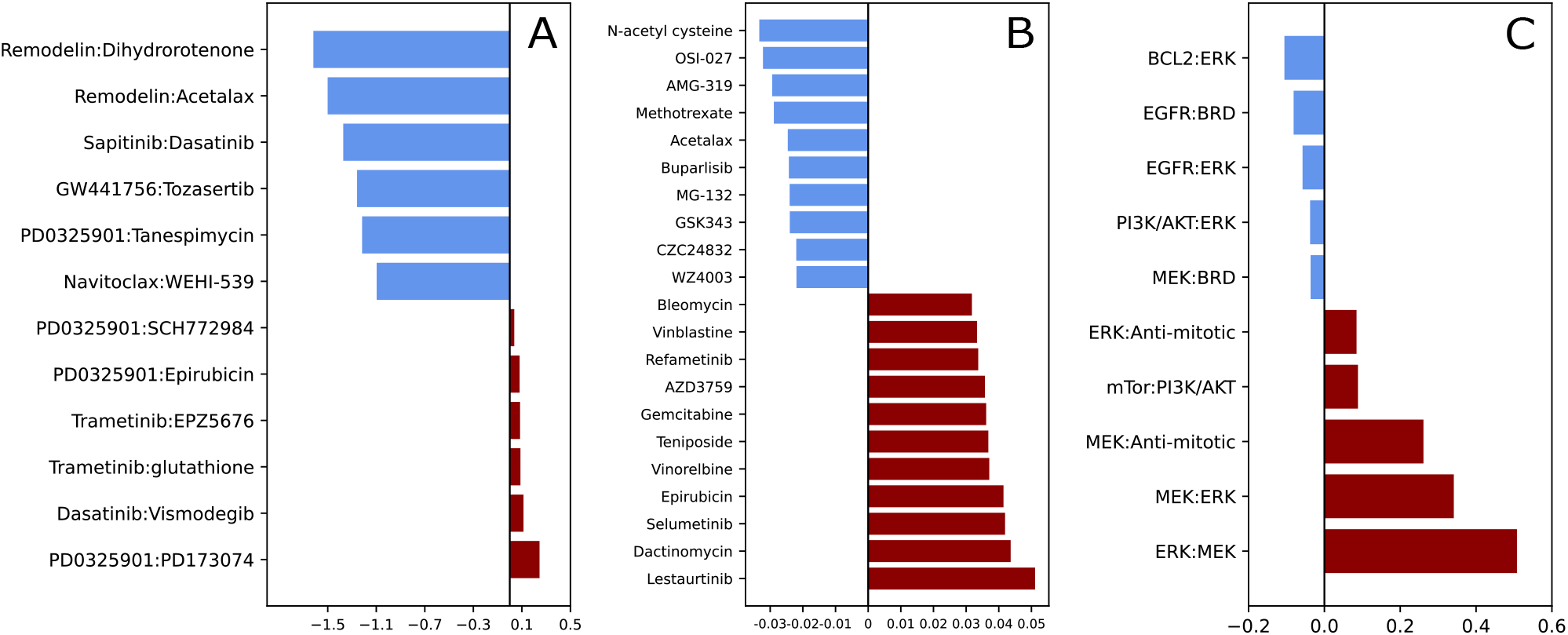
Quantification of predictive relationships across cancer drugs in drug-blind models. (A) One-to-one drug relationships (predictor:target), calculated as the logarithm of the ratio of maximum non-self predictive coefficient to self-prediction. (B) Many-to-one drug relationships, calculated as the average of the sum of all non-self predictive coefficients. (C) Class-to-class (predictor:target) relationships calculated as the average of interclass predictive coefficients.

Furthermore, mTOR inhibitors in the training set are a strong predictor for performance of PI3K and AKT inhibitors accuracy. This predictive power is stronger even than in-class relationships between mTOR inhibitors. mTOR/PI3K/AKT are strongly interrelated, so identifying where one may influence the other can be informative when making a therapeutic choice [26]. Clustering also picks out a class of anti-mitotic drugs. The anti-mitotic drug relationships are granular enough that they separate taxanes (Docetaxel and Paclitaxel) from vinca alkaloids (Vinblastine and Vinorelbine) (Fig. 4B). Anti-mitotic drugs have a class-to-class relationship with both MEK and ERK, which makes sense given the role they play in regulation of proliferation. We also can identify a family of bromodomain (BRD) inhibitors that are closely associated with one another but share almost no other predictive power with drugs outside this class. Finally, we identify a cluster of Top1/2 inhibitors. This cluster also contains Gemcitabine and Talazoparib. While both drugs do not specifically act on Top1/2, Talazoparib is a PARP inhibitor and Gemcitabine blocks DNA synthesis by mimicking deoxycytidine triphosphate. The GDSC supplied annotation identifies all of these drugs as belonging to the “DNA replication” pathway. This relationship displays that intraclass generalization refers more to a general mechanism of action than the specific target of a drug.

We identified drugs with many different predictors of performance by summing the coefficients of their predictors and averaging by the number of unique drugs in the dataset (Fig. 5B). These drugs can be identified by the vertical streaks of coefficients along the top and bottom of the heatmap in alignment with the identified drug clusters (Fig. 4A). All RAF/MEK/ERK and PI3K/AKT/mTOR drugs generalized to prediction of Lestaurtinib, a tyrosine kinase inhibitor with demonstrated target promiscuity [27]. Lestaurtinib had the strongest many-to-one relationship of all drugs in this dataset. We also saw strong many-to-one relationships for the chemotherapy drugs Bleomycin, actinomycin D, and Epirubicin. Additionally, the center of the heatmap is sparse with no strong associations outside of the diagonal. This, along with the visualization of many-to-one relationships, show there is only a small set of drugs with the ability to generalize. All of the drugs with strong potential for generalization belong to core MEK/ERK or PI3K/AKT/mTOR pathways. It is important to note that there exist no drugs with global generalization ability. These drugs, if they existed, would appear as horizontal lines in the heatmap.

More difficult to visualize in the form of a heatmap are drugs that have a specific, uniquely strong predictor of performance (Fig. 5A). We obtain one-to-one relationships by dividing the maximal drug-blind predictor of performance with the self-predictive coefficient and take the logarithm of this value. Drugs with values greater than zero are those that have a stronger predictor than themselves. There are only 6 such one-to-one relationships, with two drugs appearing across 5 of them - PD0325901 and Trametinib. Both of these drugs are MEK inhibitors. Interestingly, the drug for which PD0325901 exhibits strongest predictive power for performance is PD173074. PD173074 is a FGFR1 inhibitor, but also has been shown to inhibit the MAPK pathway [28]. Such one-to-one relationships identified from predictive performance in drug-blind models can perhaps guide experimentation routes for poorly understood drugs.

It is also important to address the presence of negative coefficients in Fig. 4. Negative coefficients indicate that a drug present in the training set makes us worse at predicting a target drug in the test set. In one-to-one and many-to-one relationships, highly negative coefficients could indicate that drugs are structurally similar but have different downstream effects. This would make them difficult to separate during embedding of Morgan fingerprints. We can see an example of this in Fig. 5B. CZC24832 and AMG-319 are both PI3K inhibitors, but they do not cluster together with other PI3K inhibitors and display some of the most negative many-to-one relationships in the whole dataset. CZC24832 specifically inhibits PI3K*γ* while AMG-319 inhibits PI3K*δ*. The isoforms of PI3K display distinctive properties, both in benign cells and in cancer, and the drugs used to specifically target them have distinct structures [29, 30]. Interestingly, other PI3K drugs do not contribute negative coefficients to AMG-319 or CZC24832. Instead, it is mostly the presence of MEK inhibitors in the training set that drive their poor drug-blind performance. Ideally, the learning of one drug should not negatively impact the learning of another. Mitigation of negative predictive power in specific situations such as the PI3K isoforms should be a goal of future work during DRP model development.

### Model training reinforces intraclass generalization in mixed set testing

We examined the embeddings created from the drug arm during mixed-set testing to determine whether intraclass similarity is increased during model training. Using the supplied pathway annotations from the GDSC2 dataset, we plotted tSNE embeddings of a subset of drugs from the families that were identified during drug-blind testing. We saw that, before training, there was some clustering of EGFR signaling and DNA replication classes when just examining raw Morgan fingerprints, but most drugs were well separated from one another (Fig. 6A). During training, we saw that by epoch 40, drugs separated into distinct clusters corresponding to mechanism (Fig. 6C). This initial separation corresponds to the steepest region of loss during training (Fig. 6B). What follows from epochs 40 to 100 appears to be more of a refinement period, where boundaries between drug mechanisms are reinforced. We observed a close relationship among MEK, EGFR, and some PI3K/mTOR drugs and another combined cluster of DNA replication and mitosis drugs. This reiterates the earlier point that while drugs with a shared mechanism are most similar in the drug feature space, models are also aware of broader pathway relationships.

**Fig 6.**
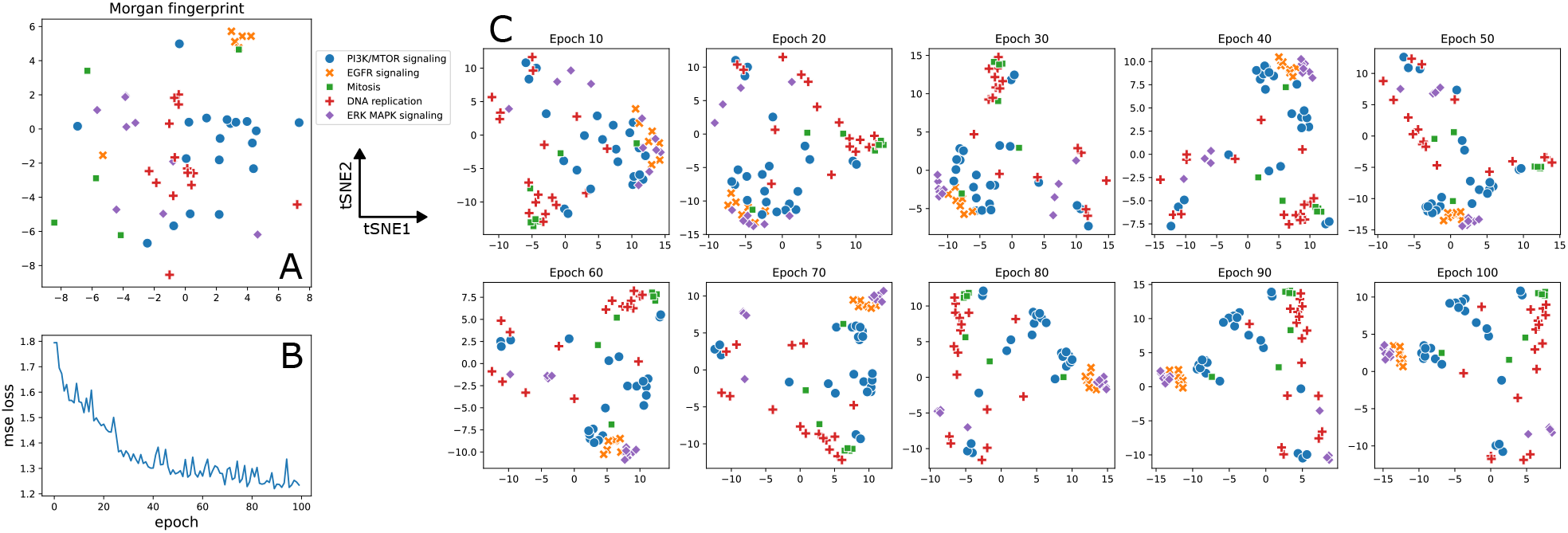
Intraclass similarities of drug embeddings increase throughout training. (A) tSNE representation of raw Morgan fingerprints for drugs with select pathways of action. There is no clear delineation between mechanisms in this representation. (B) Validation loss throughout training for drug response prediction model trained on GDSC to predict EC50. We see a steeper loss region from epochs 0 to 40 with more refinement occuring from epochs 40 to 100. (C) tSNE representation of drug embeddings at the last layer of the drug arm across 100 epochs of training. Borders between mechanisms become more defined throughout model training.

**Fig 7.**
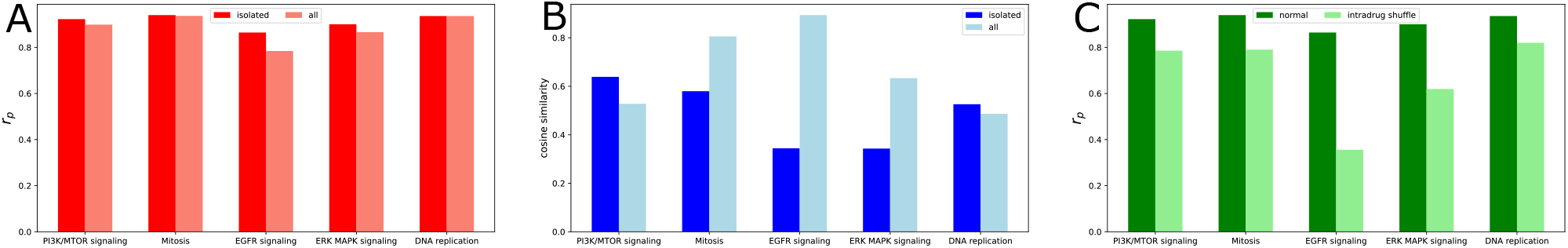
Training on specific mechanisms improves performance by decreasing intraclass similarity. Models are trained on a single mechanism (isolated) or trained on all drugs at once (all) to predict IC50 (GDSC). The test set when training on all drugs at once is created to intersect with the test set of all specific mechanisms. (A) Pearson correlation comparison in different mechanisms of action. (B) Average pairwise cosine similarity among all drugs calculated from embeddings at the last layer of the drug encoding arm of a fully trained model.

This approach also allows us to visualize how narrow the feature space occupied by each mechanism is. The features extracted by model training correspond strongly to overall mechanism of action. If a new mechanism were added to the dataset, such as in the case of a novel drug, there exists little information to correctly place it relative to all other drugs. Furthermore, there is little meaning to the space “between” clusters. Only once a mechanism is added and appropriately characterized would we be able to see the feature space it occupies.

### Training on specific drug mechanisms improves performance compared to training on all drugs at once

We sought to determine if model performance could be improved by training on only one mechanism of action at a time. We hypothesized that models trained in this manner would be better able to resolve intraclass differences in response. Training was performed on the same five drug mechanisms to be well defined during model training. Comparisons were made to a whole dataset test instance containing the exact same mechanism specific cell line and drug pairings comprising the test sets of mechanism specific models.

In all tested mechanisms, performance improves when training on a single mechanism compared to training on all mechanisms at once (Fig. 7A). We are able to achieve this performance using up to ten times less data than in whole model training. We expected that the primary source of this improvement was that the model is able to identify the specific action of individual drugs rather than collapsing all drug representations to a mechanism. Comparing the average pairwise cosine similarity of drug embeddings in each mechanism, we indeed see that drug representations become more dissimilar in the mechanism specific models that benefit most from this type of training (Fig. 7B). We then repeated the intradrug permutation experiments with these drug specific models (Fig. 7C). Mechanism specific training made two of our studied pathways more sensitive to intradrug permutation, with EGFR being five times more sensitive than the baselines seen in 1. Again, this occurs by the largest margin in mechanisms for whom mechanism specific training is the most beneficial to performance. In these models, drugs require the correct cell line information to remain accurate. Decreased cosine similarity and improved intradrug permutation sensitivity are not present in in PI3K/mTOR and DNA replication specific models. This is likely due to the fact that these classes contain drugs with more variety in their specific targets.

## Discussion

Over 30 years ago, the NCI60 team displayed that a neural network had the ability to predict drug mechanism of action based solely on cell line responses [31]. Even given advanced representations of cell lines and expansion of pharmacogenomic datasets to hundreds of thousands of experiments, cancer drug response prediction models still only collapse to representing mechanism of action in order to infer responses. Drug-blind failure simply describes something that has already been shown: cell line based drug response can be simplified to a function of drug mechanism of action. Drug-blind testing, then, should not be viewed as an appropriate benchmark for cancer drug response prediction models. Examining the behavior of models in a mechanism specific context is more pertinent to understanding drug generalization in the context of large pharmacogenomic datasets. This would involve training and testing models according to more specific use cases, such as a particular patient populations.

Drug-blind prediction is primarily relevant to identifying the efficacy of novel compounds. An ideal model with a global understanding of cancer drug generalizability would be adept at determining the effectiveness of any given drug in a zero-shot setting, not just one for whom it understands the mechanism of action. We found that we cannot expect this global generalizability in cancer drugs in the context of currently existing datasets and drugs, but there is still utility for examining understudied compounds. 63% of approvals of compounds for cancer treatment by the FDA from 2009 to 2020 were next-in-class, i.e. they used a previously described mechanism of action [32]. If drug-blind prediction is capable of intraclass generalization, then it is capable of generalizing to prediction of performance for the majority of newly approved compounds. In fact, mixed set training behaves inherently like contrastive learning among the known mechanisms of drugs (Fig. 6C).

Supervised contrastive learning can be used to train models that amplify features of intraclass similarity [33, 34]. Cancer DRP models recreate this behavior without any class labels or high similarity of Morgan fingerprints, only by learning shared cell lines responses. Next-in-class prediction is therefore a realistic drug-blind task, especially with models tuned to specific mechanisms. A potential solution for prediction within new mechanisms of action would be to create models with built in uncertainty, such as in [35]. Given the sparsity of generalizability, drug-blind models should be aware of what they do not know given the information they already have to prevent inaccurate predictions. Drug-blind models could then mark drugs as poorly understood or even for further experimentation, such as in a lab automation setting.

There is an argument to be made that some representations of drugs in deep learning could be more generalizable than others. That is, these representations would better bridge the feature space among drug mechanisms of action. Because this paper is specifically intended to be data-centric, we use a constant MLP architecture and Morgan fingerprint to probe dataset behavior. Among other studies that examine drug-blind prediction, drugs are represented using molecular graphs and graph neural networks [36–39], word embeddings [19, 40], and canonical SMILES for convolution [41]. We refer readers to [4] for a more complete review of studies that utilize drug-blind benchmarks. Of the results that predict continuous drug response values and report pearson correlation for comparison, none outperforms the distribution we observe in this paper (Fig. 3). Drug-blind failure is a global phenomenom, not architecture specific. The results of this paper also suggest that given the wide distribution of performance values, it is essential that an appropriately large amount of training splits are tested. This is because drug-blind analysis is so sensitive to the chosen train/test split due to changing mechanistic overlap. Only [41] performs a more appropriate amount of replicates, with 150, while others perform 3- and 5-fold cross validation. We recommend that future attempts to benchmark any improvement in drug-blind performance analyze a sufficient amount of replicates to cover the number of unique drugs in the dataset. Furthermore, it appears that optimization based approaches trump architectural design when considering approaches to directly improve drug-blind generalizability [21]. In other words, changing how drugs share information is more important than changing what information is shared. As we observe in this paper, the amount of information that drugs across mechanistic classes are able to share is quite minimal regardless.

When identifying treatments for patients, physicians often determine candidate drugs using a particular mechanism of action that addresses a concrete event, such as a mutation. This, in theory, translates better to a classification task where we can determine whether a certain drug exhibits the desired mechanism of action. As we saw, this is a task DRP models are highly capable of. Unfortunately, empirically determining mechanism of action is a difficult task and results often do not belong to one discrete class [27, 42]. As we have seen, the majority of drugs do not cluster into a well defined class in a drug-blind setting (Fig. 4). Take, for example, the case of Lestaurtinib both in this paper and in [27]. The advertised target kinase of Lestaurtinib is not even in the top 20 targets for which it is actually the most active. Indeed, Klaeger et. al find that this is the case for many different kinase inhibitors. In the context of this promiscuous behavior of drugs, how do we identify effectiveness given particular cancer cell behavior? The answer to this question lies in the interpretation of many-to-one relationships in drug-blind prediction. The results in Fig. 5B should not necessarily suggest that Lestaurtinib can function as a MEK, EGFR, or ERK inhibitor. Rather, the downstream effects of Lestaurtinib and these core pathways merge at some point. Since we know that Lestaurtinib can exhibit such a broad array of bindings, it is more likely for its inhibitory potential to overlap with another given drug’s at any time. For example, it has been found that Lestaurtinib exhibits synergistic effects through MAST1 mediated MEK activation [43].

Generalization outside of a specific mechanism of action is perhaps then more indicative of potential drug synergy. Recent work has shown that categorical embeddings of drug mechanisms can be used to enhance drug synergy prediction [44]. Rather than training categorical embeddings directly, embeddings from DRP models could perhaps be used as they encode mechanistic information while maintaining flexibility as in the case of Lestaurtinib. Drug synergy prediction is an entirely different, and potentially more clinically relevant, task in cancer DRP [45]. We are not recommending that we only perform classification of mechanism in drugs, but rather that our approach to analysis of drug-blind testing can better describe these drugs with more ambiguous activity. Determining drug-to-drug performance benefit is perhaps more helpful for understanding ambiguous drugs’ responses than predicting some continuous response variable itself.

## Conclusion

In an ideal dataset, there would be enough drugs to cover the entire feature space that spans all mechanisms of action and therefore allows for maximum generalizability between drugs. This is the logic behind the suggestion that the drop in drug blind performance is simply because we do not have datasets with enough cancer drugs. Assuming that there is some point at which we will have collected enough data from cell lines and drugs such that we have a complete and generalizable understanding of the cancer drug feature space is a distant goal.

Instead, generation of mechanism focused data and designing specialized DRP models holds great promise for an improved understanding of cancer biology through AI.

## Materials and methods

### Drug response datasets

All cell lines were represented using gene expression data. Expression data was obtained from DepMap using Expression Public 24Q2 [46]. Cell lines without expression data from DepMap were filtered out from their respective datasets. We used DepMap as a constant source for gene expression data due to potential experimental differences across other collections and need to minimize sources of variance in model input. Gene expression features were filtered to intersect with patient expression data from the Cancer Genome Atlas (TCGA) [47].

Drugs were represented as Morgan fingerprints for input to our model. We first obtained the SMILES string for each drug using the PubChemPy python library. The PubChem database was queried using the supplied name of the drug from each dataset and the best match was used. Morgan fingerprints were generated using the GetMorganGenerator function of RDKit with a radius size of 2 to return a fingerprint vector of length 2048.

Drug SMILES were not manually annotated. If they were not automatically retrieved from PubChem, they were removed from the dataset. Reported dataset sizes below are after removal of cell lines without available gene expression data and drugs without an available PubChem SMILES.

1. **gCSI [48]** The Genentech Cell Line Screening Initiative was obtained from http://research-pub.gene.com/gCSI_GRvalues2019/. This dataset was of particular interest as it includes growth rate metrics (GR50, etc.) as an alternative to IC50. There were 43 unique drugs. Number of unique cell lines varied by metric based on available data.
2. **CTRPv2 [49, 50]** Data from the Cancer Therapeutics Response Portal (v2) was obtained from the DepMap portal under the CTD^2 release. There were 496 unique drugs and 839 unique cell lines present in the dataset.
3. **GDSC2 [51, 52]** Data from Genomics for Drug Sensitivity were downloaded from https://www.cancerrxgene.org/downloads/bulk_download. We specifically utilized GDSC2. There were 230 unique drugs and 936 unique cell lines present in the dataset.

Initial permutation experiments were performed across all three datasets and all related metrics. All further experiments were performed on only GDSC2 using EC50. The code used for all data preprocessing and experiments in this study is provided at https://github.com/willherbert27/drug_blind_generalization.

### Model architecture and optimization

The deep learning model architecture used in this work is comprised of a multilayer perceptron (MLP) with two arms for separate embedding of drugs and cell lines (Fig. 8). Cell line and drug representations are then concatenated and used to predict a continuous response value defined by the input dataset. Input to the model was therefore in the form of (cell line, drug, response) tuples. This simple model architecture was chosen in order to provide a constant baseline that was able to train rapidly with good performance. This architecture is similar to MLPs used for DRP dataset examination in prior work [8, 9, 53]. Unless otherwise specified, models were trained with a batch size of 16 using the Adam optimizer with a learning rate of 1e-4. Models were trained using early stopping. A validation set was used to determine the epoch of early stopping and all performance is reported using a further test set. Data was divided into train/validation/test sets using an 80/10/10 splitting strategy. It should be reiterated that the primary focus of this work was not to spend time identifying a hyperparameter tuning strategy to maximize performance under a specific architecture. Instead, we maintain a simple and constant model training strategy in order to more clearly examine the impact of data the model is trained on.

**Fig 8.**
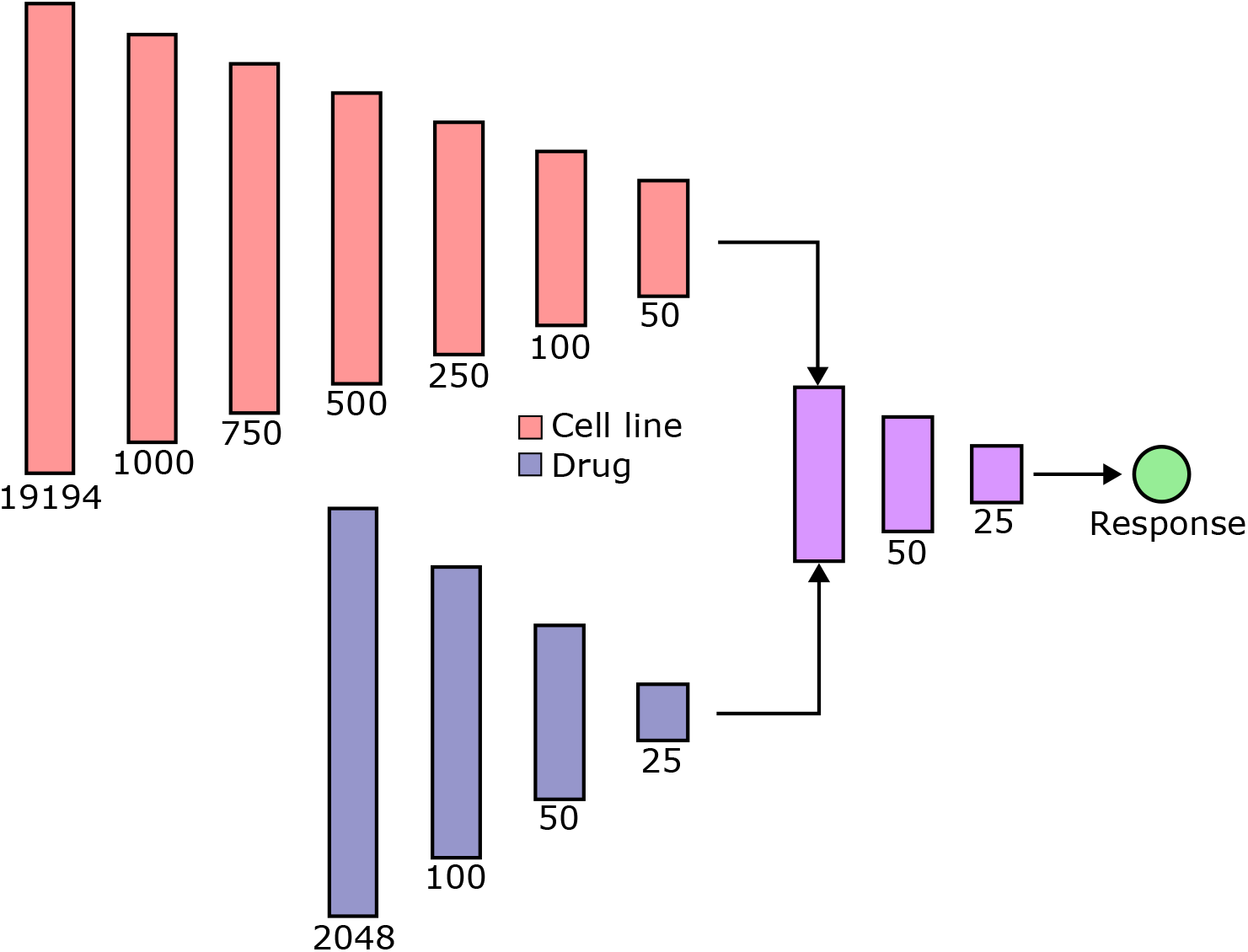
Model architecture. The model used in this study is comprised of a multilayer perceptron with separate drug embedding (blue) and cell line embedding (red) arms. These embeddings are then concatenated to calculate a given response. All fully connected layers, excluding output, are followed by ReLU activation.

### Permutation experiments

Intradrug model permutation was performed by randomly shuffling response values across all cell lines to which a particular drug was applied, as in [17], i.e. for a single drug, the distribution of response values went unchanged. The response values were assigned to a different cell line than the original. In theory, this should disconnect drug representations as a predictor of its response from using specific information about a cell line it is applied to. We also randomly permuted values across all drugs applied to a particular cell line, called intracell shuffling. In one-hot encoding, cell lines were represented with one-hot vectors rather than gene expression.

### Diversity of dataset composition

Dataset composition experiments were performed using the GDSC dataset. When testing the impact of the number of cell lines present for a given drug, the dataset was partitioned to only include drugs tested on 500 or more cell lines. This allowed the pool of drugs to remain the same across all diversity experiments. Filtering the drugs in this way reduced unique drugs from 230 to 214 drugs in GDSC2. The training set was then further filtered to only include the specified number of cell lines. Drug-blind experiments refer to those for which the drugs available in the training, validation, and test sets are disjoint from one another. When examining the impact of quantity of unique cell lines on mixed-set performance, the test set was held constant across all replicates. This was done in order to have a constant set for quantification of increased generalization. Five replicates were performed for each experiment in order to report standard deviation.

### Prediction of performance by training set composition

By varying the data present in the training set of a model, we sought to determine the impact of the inclusion of different drugs on model performance. For these experiments, we again utilize the GDSC2 dataset. The test and validation sets across all experiments are held constant at 10% (23) of unique drugs while the training set is varied. The total number of drugs in the training set at any given time was comprised of 50% (115) of the total unique drugs. To predict drug-blind performance based on training set composition, we trained an elastic net model [54] using scikit-learn (alpha=0.01,lambda=0.1) using training set composition as features and accuracy as targets. Features were binary vectors indicating the presence or absence of a particular drug in the training set and target values were the calculated Pearson correlation of that particular experiment.

To further elucidate the relationship between individual drugs, we created an experimental setup where the composition of drugs in the training set varies but the test set contains all 230 unique drugs in the dataset. In this way, the test set contains both drug-blind and non-drug-blind examples. We do this so that we have a constant set of targets with which to fit linear models after DRP model training. The training set was again partitioned to include 50% (115) of the unique drugs present in the dataset. The validation and test sets contained the remaining non-overlapping cell lines for drugs present in the training set as well as all examples for drug-blind pairs. We trained 1,641 drug response prediction models total for these experiments. We then calculated the performance on a drug-by-drug basis for each DRP model using Pearson correlation. Because we can obtain individual drug accuracy for all drugs across all models, we can fit an elastic net model to the performance of each drug. Input features are the drugs present in the training set, binarized, and targets are per model pearson correlation for the current drug. We summed and averaged the coefficients obtained for each drug across 10-fold cross validation. Using these coefficients, we performed hierarchical clustering on the predictive ability of each drug in the training set.

## Acknowledgments

We would like to thank Esther Rodman for her helpful discussions on cancer drug mechanisms of action. We thank members of the IMPROVE project for allowing us to join in on regular meetings and helping us to build the knowledge base necessary for this work. We would also like to thank all members of the Walther-Antonio laboratory for their feedback during project development and writing.

## References

1. Siegel RL, Miller KD, Jemal A. Cancer statistics, 2018. CA: a cancer journal for clinicians. 2018;68(1):7–30.

2. Swanton C, Bernard E, Abbosh C, André F, Auwerx J, Balmain A, et al. Embracing cancer complexity: Hallmarks of systemic disease. Cell. 2024;187(7):1589–1616.

3. Baptista D, Ferreira PG, Rocha M. Deep learning for drug response prediction in cancer. Briefings in bioinformatics. 2021;22(1):360–379.

4. Partin A, Brettin TS, Zhu Y, Narykov O, Clyde A, Overbeek J, et al. Deep learning methods for drug response prediction in cancer: predominant and emerging trends. Frontiers in medicine. 2023;10:1086097.

5. Partin A, Vasanthakumari P, Narykov O, Wilke A, Koussa N, Jones SE, et al. Benchmarking community drug response prediction models: datasets, models, tools, and metrics for cross-dataset generalization analysis. arXiv preprint arXiv:250314356. 2025;.

6. Overbeek JC, Partin A, Brettin TS, Chia N, Narykov O, Vasanthakumari P, et al. Assessing Reusability of Deep Learning-Based Monotherapy Drug Response Prediction Models Trained with Omics Data. arXiv preprint arXiv:240912215. 2024;.

7. Piyawajanusorn C, Nguyen LC, Ghislat G, Ballester PJ. A gentle introduction to understanding preclinical data for cancer pharmaco-omic modeling. Briefings in Bioinformatics. 2021;22(6):bbab312.

8. Partin A, Brettin T, Evrard YA, Zhu Y, Yoo H, Xia F, et al. Learning curves for drug response prediction in cancer cell lines. BMC bioinformatics. 2021;22:1–18.

9. Xia F, Allen J, Balaprakash P, Brettin T, Garcia-Cardona C, Clyde A, et al. A cross-study analysis of drug response prediction in cancer cell lines. Briefings in bioinformatics. 2022;23(1):bbab356.

10. Gillet JP, Varma S, Gottesman MM. The clinical relevance of cancer cell lines. Journal of the National Cancer Institute. 2013;105(7):452–458.

11. Raji ID, Kumar IE, Horowitz A, Selbst A. The fallacy of AI functionality. In: Proceedings of the 2022 ACM Conference on Fairness, Accountability, and Transparency; 2022. p. 959–972.

12. Sambasivan N, Kapania S, Highfill H, Akrong D, Paritosh P, Aroyo LM. “Everyone wants to do the model work, not the data work”: Data Cascades in High-Stakes AI. In: proceedings of the 2021 CHI Conference on Human Factors in Computing Systems; 2021. p. 1–15.

13. Haibe-Kains B, El-Hachem N, Birkbak NJ, Jin AC, Beck AH, Aerts HJ, et al. Inconsistency in large pharmacogenomic studies. Nature. 2013;504(7480):389–393.

14. Lenhof K, Eckhart L, Rolli LM, Lenhof HP. Trust me if you can: a survey on reliability and interpretability of machine learning approaches for drug sensitivity prediction in cancer. Briefings in Bioinformatics. 2024;25(5):bbae379.

15. Safikhani Z, Smirnov P, Freeman M, El-Hachem N, She A, Rene Q, et al. Revisiting inconsistency in large pharmacogenomic studies. F1000Research. 2017;5:2333.

16. Gura T. Systems for identifying new drugs are often faulty; 1997.

17. Yuan H, Paskov I, Paskov H, González AJ, Leslie CS. Multitask learning improves prediction of cancer drug sensitivity. Scientific reports. 2016;6(1):31619.

18. Park A, Lee Y, Nam S. A performance evaluation of drug response prediction models for individual drugs. Scientific Reports. 2023;13(1):11911.

19. Branson N, Cutillas P, Bessant C. Understanding the Sources of Performance in Deep Learning Drug Response Prediction Models. bioRxiv. 2024;.

20. Codicè F, Pancotti C, Rollo C, Moreau Y, Fariselli P, Raimondi D. The specification game: rethinking the evaluation of drug response prediction for precision oncology. Journal of Cheminformatics. 2025;17(1):33.

21. Narykov O, Zhu Y, Brettin T, Evrard YA, Partin A, Xia F, et al. Data imbalance in drug response prediction: multi-objective optimization approach in deep learning setting. Briefings in Bioinformatics. 2025;26(2):bbaf134.

22. Tan J, Yang J, Wu S, Chen G, Zhao J. A critical look at the current train/test split in machine learning. arXiv preprint arXiv:210604525. 2021;.

23. Martin TM, Harten P, Young DM, Muratov EN, Golbraikh A, Zhu H, et al. Does rational selection of training and test sets improve the outcome of QSAR modeling? Journal of chemical information and modeling. 2012;52(10):2570–2578.

24. Chappell WH, Steelman LS, Long JM, Kempf RC, Abrams SL, Franklin RA, et al. Ras/Raf/MEK/ERK and PI3K/PTEN/Akt/mTOR inhibitors: rationale and importance to inhibiting these pathways in human health. Oncotarget. 2011;2(3):135.

25. McCubrey JA, Steelman LS, Chappell WH, Abrams SL, Montalto G, Cervello M, et al. Mutations and deregulation of Ras/Raf/MEK/ERK and PI3K/PTEN/Akt/mTOR cascades which alter therapy response. Oncotarget. 2012;3(9):954.

26. Dienstmann R, Rodon J, Serra V, Tabernero J. Picking the point of inhibition: a comparative review of PI3K/AKT/mTOR pathway inhibitors. Molecular cancer therapeutics. 2014;13(5):1021–1031.

27. Klaeger S, Heinzlmeir S, Wilhelm M, Polzer H, Vick B, Koenig PA, et al. The target landscape of clinical kinase drugs. Science. 2017;358(6367):eaan4368.

28. Nguyen P, Tsunematsu T, Yanagisawa S, Kudo Y, Miyauchi M, Kamata N, et al. The FGFR1 inhibitor PD173074 induces mesenchymal–epithelial transition through the transcription factor AP-1. British journal of cancer. 2013;109(8):2248–2258.

29. Zhao L, Vogt PK. Class I PI3K in oncogenic cellular transformation. Oncogene. 2008;27(41):5486–5496.

30. Vanhaesebroeck B, Perry MW, Brown JR, André F, Okkenhaug K. PI3K inhibitors are finally coming of age. Nature reviews Drug discovery. 2021;20(10):741–769.

31. Weinstein JN, Kohn KW, Grever MR, Viswanadhan VN, Rubinstein LV, Monks AP, et al. Neural computing in cancer drug development: predicting mechanism of action. Science. 1992;258(5081):447–451.

32. Olivier T, Haslam A, Prasad V. Anticancer drugs approved by the US Food and Drug Administration from 2009 to 2020 according to their mechanism of action. JAMA network open. 2021;4(12):e2138793–e2138793.

33. Khosla P, Teterwak P, Wang C, Sarna A, Tian Y, Isola P, et al. Supervised contrastive learning. Advances in neural information processing systems. 2020;33:18661–18673.

34. Graf F, Hofer C, Niethammer M, Kwitt R. Dissecting supervised contrastive learning. In: International Conference on Machine Learning. PMLR; 2021. p. 3821–3830.

35. Lenhof K, Eckhart L, Rolli LM, Volkamer A, Lenhof HP. Reliable anti-cancer drug sensitivity prediction and prioritization. Scientific Reports. 2024;14(1):12303.

36. Zhu Y, Ouyang Z, Chen W, Feng R, Chen DZ, Cao J, et al. TGSA: protein–protein association-based twin graph neural networks for drug response prediction with similarity augmentation. Bioinformatics. 2022;38(2):461–468.

37. Wang C, Kumar GA, Rajapakse JC. Drug discovery and mechanism prediction with explainable graph neural networks. Scientific Reports. 2025;15(1):179.

38. Nguyen T, Nguyen GT, Nguyen T, Le DH. Graph convolutional networks for drug response prediction. IEEE/ACM transactions on computational biology and bioinformatics. 2021;19(1):146–154.

39. Liu Q, Hu Z, Jiang R, Zhou M. DeepCDR: a hybrid graph convolutional network for predicting cancer drug response. Bioinformatics. 2020;36(Supplement 2):i911–i918.

40. Jiang L, Jiang C, Yu X, Fu R, Jin S, Liu X. DeepTTA: a transformer-based model for predicting cancer drug response. Briefings in bioinformatics. 2022;23(3):bbac100.

41. Liu P, Li H, Li S, Leung KS. Improving prediction of phenotypic drug response on cancer cell lines using deep convolutional network. BMC bioinformatics. 2019;20:1–14.

42. Hight SK, Clark TN, Kurita KL, McMillan EA, Bray W, Shaikh AF, et al. High-throughput functional annotation of natural products by integrated activity profiling. Proceedings of the National Academy of Sciences. 2022;119(49):e2208458119.

43. Jin L, Chun J, Pan C, Li D, Lin R, Alesi GN, et al. MAST1 drives cisplatin resistance in human cancers by rewiring cRaf-independent MEK activation. Cancer cell. 2018;34(2):315–330.

44. Gonzàlez Lastre M, Gonzàlez De Prado Salas P, Guantes R. Optimizing drug synergy prediction through categorical embeddings in Deep Neural Networks. Biology Methods and Protocols. 2025;10(1):bpaf033.

45. Torkamannia A, Omidi Y, Ferdousi R. A review of machine learning approaches for drug synergy prediction in cancer. Briefings in Bioinformatics. 2022;23(3):bbac075.

46. DepMap B. DepMap 24Q2 Public; 2024. Available from: 10.25452/figshare.plus.27993248.v1.

47. Weinstein JN, Collisson EA, Mills GB, Shaw KR, Ozenberger BA, Ellrott K, et al. The cancer genome atlas pan-cancer analysis project. Nature genetics. 2013;45(10):1113–1120.

48. Haverty PM, Lin E, Tan J, Yu Y, Lam B, Lianoglou S, et al. Reproducible pharmacogenomic profiling of cancer cell line panels. Nature. 2016;533(7603):333–337.

49. Rees MG, Seashore-Ludlow B, Cheah JH, Adams DJ, Price EV, Gill S, et al. Correlating chemical sensitivity and basal gene expression reveals mechanism of action. Nature chemical biology. 2016;12(2):109–116.

50. Seashore-Ludlow B, Rees MG, Cheah JH, Cokol M, Price EV, Coletti ME, et al. Harnessing connectivity in a large-scale small-molecule sensitivity dataset. Cancer discovery. 2015;5(11):1210–1223.

51. Yang W, Soares J, Greninger P, Edelman EJ, Lightfoot H, Forbes S, et al. Genomics of Drug Sensitivity in Cancer (GDSC): a resource for therapeutic biomarker discovery in cancer cells. Nucleic acids research. 2012;41(D1):D955–D961.

52. Iorio F, Knijnenburg TA, Vis DJ, Bignell GR, Menden MP, Schubert M, et al. A landscape of pharmacogenomic interactions in cancer. Cell. 2016;166(3):740–754.

53. Xia F, Shukla M, Brettin T, Garcia-Cardona C, Cohn J, Allen JE, et al. Predicting tumor cell line response to drug pairs with deep learning. BMC bioinformatics. 2018;19:71–79.

54. Zou H, Hastie T. Regularization and variable selection via the elastic net. Journal of the Royal Statistical Society Series B: Statistical Methodology. 2005;67(2):301–320.

